# Deep learning-based prediction of chemical accumulation in a pathogenic mycobacterium

**DOI:** 10.1101/2024.12.15.628588

**Authors:** Mark R. Sullivan, Eric J. Rubin

## Abstract

Drugs must accumulate at their target site to be effective, and inadequate uptake of drugs is a substantial barrier to the design of potent therapies. This is particularly true in the development of antibiotics, as bacteria possess numerous barriers to prevent chemical uptake. Designing compounds that circumvent bacterial barriers and accumulate to high levels in cells could dramatically improve the success rate of antibiotic candidates. However, a comprehensive understanding of which chemical structures promote or prevent drug uptake is currently lacking. Here we use liquid chromatography-mass spectrometry to measure accumulation of 1528 approved drugs in *Mycobacterium abscessus*, a highly drug-resistant, opportunistic pathogen. We find that simple chemical properties fail to effectively predict drug accumulation in mycobacteria. Instead, we use our data to train deep learning models that predict drug accumulation in *M. abscessus* with high accuracy, including for chemically diverse compounds not included in our original drug library. We find that differential drug uptake is a critical determinant of the efficacy of drugs currently in development and can identify compounds which accumulate well and have antibacterial activity in *M. abscessus*. These predictive algorithms can be an important complement to chemical synthesis and accumulation assays in the evaluation of drug candidates.

## Introduction

Drug accumulation and retention has long been seen as a barrier to effective therapy, from cancer treatments to antibiotic regimens^1–3^. Drug accumulation is driven by a complex combination of entry, efflux, and metabolism within cells. In bacteria, all three of these processes present substantial hurdles to drug retention. Antibiotics first must pass through the cell envelope, which varies dramatically in structure across genera^4,5^, and therefore likely excludes different types of compounds. Once a chemical has entered a bacterium, it can be subject to active efflux^2,6^. Efflux pumps are present across most bacterial species, and have a wide range of substrate specificities that include many naturally occurring antibiotics^2,6^. Chemicals that pass those hurdles must also avoid being metabolized by the cell as bacteria possess numerous enzymes the enact chemical modification and inactivation of drugs^7^. These intrinsic barriers to drug accumulation vary in strength across bacterial species; some species are generally susceptible to drugs, while others possess multiple barriers to chemical accumulation and are therefore highly resistant to many antibiotics. Bacteria that effectively exclude drugs in this way can be clinically challenging pathogens. One such organism is *Mycobacterium abscessus*. *M. abscessus* is an opportunistic systemic and pulmonary pathogen that tends to infect individuals with predisposing conditions like T cell defects, cystic fibrosis and chronic obstructive pulmonary disease^8,9^. Due to this organism’s broad-spectrum, intrinsic resistance to antibiotics, treatments often fail, with cures achieved only 30-50% of the time^10–13^. Identifying chemical structures that are able to effectively evade the barrier functions of bacteria like *M. abscessus* and accumulate to toxic levels could enable more rapid development of effective therapies.

Numerous efforts have been made to identify determinants of drug uptake in bacteria^6,14–17^. In the Gram-negative bacteria *Escherichia coli* and *Pseudomonas aeruginosa*, drug accumulation into cells is more effective for positively charged compounds compared to uncharged chemicals^14,16^, and rigid, non-globular compounds accumulate most readily in *E. coli*^16^. Further, chemical labeling methods in mycobacteria that have enabled the measurement of chemical penetration across the cell envelope but not into the cell suggest that the cell envelope represents a critical, selective barrier to chemical accumulation^15^. These insights into drug accumulation demonstrate that there are rational chemical and biological rules that dictate whether a compound will be retained in a particular bacterium. However, our current ability to identify those rules is largely limited to general chemical properties that likely do not capture the full range of characteristics that determine whether a compound will accumulate.

Here, to uncover the chemical rules that dictate drug accumulation in mycobacteria, we use liquid chromatography-mass spectrometry (LC-MS) to measure the accumulation in *M. abscessus* of a chemically diverse library of 1528 approved drugs. We then examine whether physical properties of chemicals predict accumulation in *M. abscessus*. Finally, we develop a deep learning-based model that accurately predicts chemical accumulation in *M. abscessus* based directly on chemical structure information.

## Results

### Approved drugs display a wide range of accumulation in *M. abscessus*

To understand the rules that dictate chemical uptake, we sought to measure accumulation of a chemically diverse array of compounds in *M. abscessus*. Since drug-like compounds are the chemicals most representative of future antibiotic candidates, we tested whether a library of 2320 drugs (APExBIO DiscoveryProbe FDA-approved Drug Library) that have been previously approved for use in humans or animals could be measured by a single LC-MS method. 1528 of these compounds could be detected and quantitated over a linear range (Supplemental File 1), enabling the measurement of accumulation for these compounds. To quantitate drug buildup in bacteria, we adapted a previous method^18^ to measure accumulation of many compounds in parallel in *M. abscessus* (Figure 1A). Accumulation of these compounds spanned a range of greater than eight orders of magnitude and was robust across biological replicates (Figure 1B) as well as between duplicate versions of compounds with different counter-ions or protonation states (Figure 1C). Together, these results suggest that multiplexed LC-MS provides a robust, repeatable measurement of drug accumulation.

**Figure 1.**
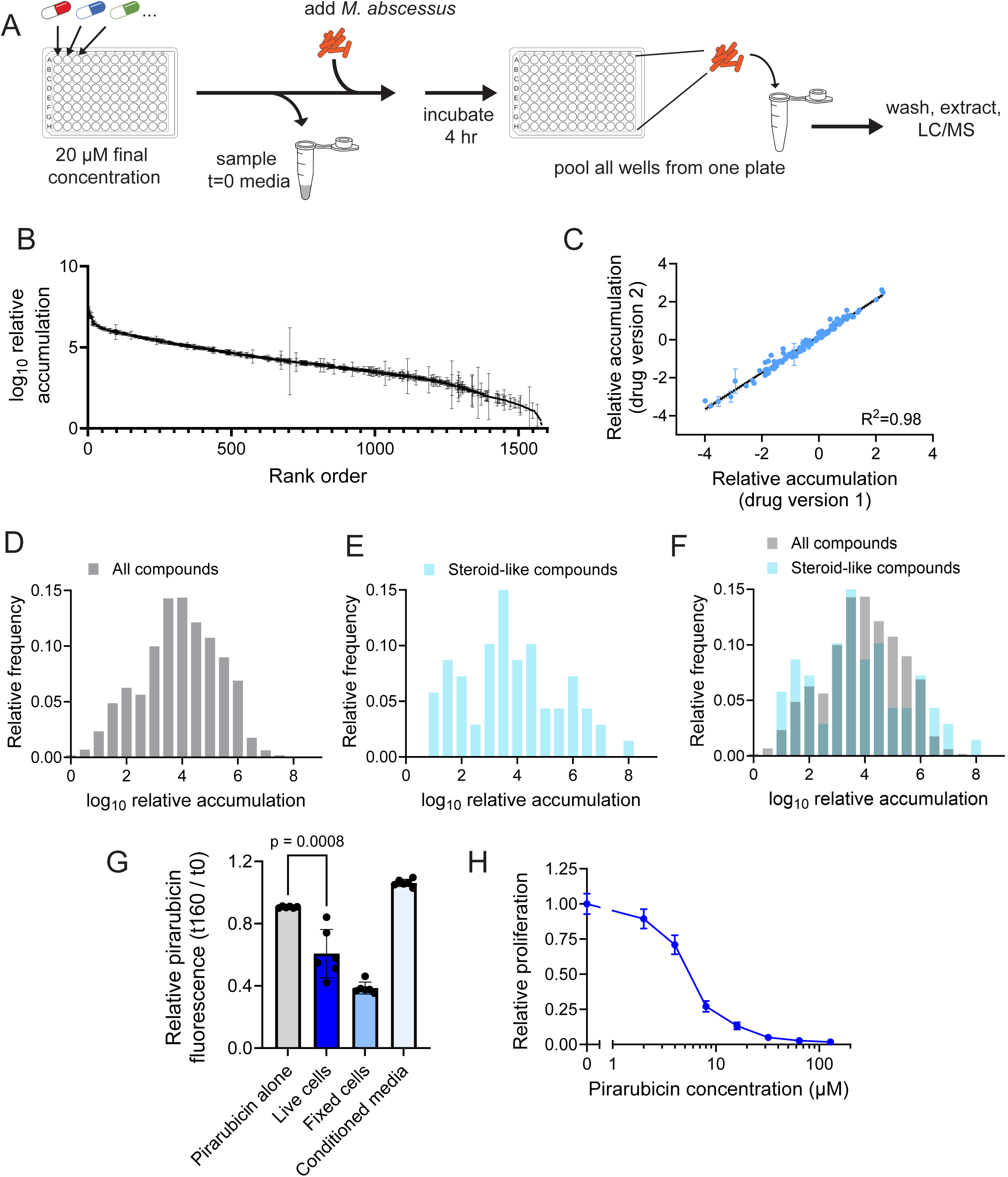
Measurement of accumulation of approved drugs in *M. abscessus*. **(A)** Schematic of method for measurement of accumulation of library of 1528 compounds approved for use in humans or animals in *M. abscessus*. LC-MS: liquid chromatography-mass spectrometry. **(B)** LC-MS measurement of peak area of cell-associated fraction of 1528 compounds in *M. abscessus* normalized to peak area of each compound in the media used for the experiment prior to incubation with bacteria. Relative accumulation is normalized to the compound with the lowest accumulation. Data represent mean +/- SD. n=3 independent cultures for each chemical compound. **(C)** Correlation between relative accumulation of compounds replicated in the chemical library with different counter-ions or protonation state. Each dot represents mean +/- SD of one compound. n=3 independent cultures for each compound. **(D)** Histogram displaying the relative accumulation of all 1528 compounds **(D)**, only steroid-like compounds **(E)**, or both overlaid **(F)** in *M. abscessus*. Relative accumulation was calculated as described in **(B)**. **(F)** Fluorescence of pirarubicin after 160 minutes of incubation normalized to initial fluorescence. Fluorescence was measured either after incubation of pirarubicin in fresh culture medium (pirarubicin alone), conditioned culture medium, with live *M. abscessus*, or with fixed *M. abscessus*. Data represent mean +/- SD. n=6 independent cultures for each condition. p-value derived from unpaired, two-tailed t test. **(G)** Relative proliferation of *M. abscessus* as measured by autoluminescence in the presence of the indicated concentrations of pirarubicin. Proliferation is normalized to vehicle-treated condition. Data represent mean +/- SD. n=6 independent cultures for each drug dose.

Chemical accumulation is likely driven by uptake, not non-specific binding These measurements of drug accumulation could be driven either by chemicals entering into cells or by being bound tightly enough to the exterior of cells to not become dislodged during washing. Given the lipid-rich nature of the mycobacterial envelope, the drugs most likely to adhere non-specifically to the outside of the cell are lipophilic compounds. As a result, if non-specific binding of compounds to *M. abscessus* is driving accumulation, we would predict that lipophilic classes of compounds would universally display high accumulation. One of the most well-represented classes of compound in the library used for LC-MS analysis are steroids, which are highly lipophilic and capable of being incorporated into membranes. Thus, if accumulation measurements represent non-specific binding to the cell envelope, these lipophilic steroids should all display high accumulation. Instead, though some steroids do accumulate well, the overall distribution of steroid-like compounds is similar to that of total compounds (Figure 1D-F, Supplemental Figure 1A). This suggests that factors other than non-specific binding to the cell membrane are responsible for the distribution of chemical accumulation values.

To further test whether compounds with high measured accumulation enter the bacterium, we made use of the natural fluorescence of pirarubicin, a drug that was in the top 1% of accumulators. Pirarubicin is a naturally fluorescent DNA intercalator, and its fluorescence is quenched once it binds DNA^19^. Thus, a decrease in pirarubicin fluorescence indicates that it has entered the cell and bound to DNA, rather than adhered to the outside of the bacterium. Incubation of pirarubicin with fixed, permeabilized *M. abscessus* leads to a decrease in pirarubicin fluorescence (Figure 1G). Incubation with live *M. abscessus* also results in a substantial reduction in fluorescence, demonstrating that pirarubicin accumulates within live bacterial cells (Figure 1G). This fluorescence quenching is not a result of extracellular DNA-binding, as incubation of pirarubicin in conditioned media from *M. abscessus* does not result in decreased fluorescence (Figure 1G). Though pirarubicin is a cancer drug^20^, it also displays antibacterial activity^21^, so we reasoned that if it is taken up to a high degree, it may be toxic to *M. abscessus*. Consistent with this hypothesis, pirarubicin prevents bacterial growth (Figure 1H) at doses comparable to antibiotics clinically used to treat *M. abscessus*^22^. Together, these results suggest that non-specific binding to the bacterium is not a major determinant of drug accumulation and that the most highly accumulating compounds are likely entering cells at levels sufficient to exhibit biological effects.

### Antibiotics accumulate poorly in *M. abscessus*

We next surveyed patterns of chemical accumulation in *M. abscessus*. Given that the drug library contained numerous antimicrobial compounds, we first examined whether any display effective accumulation in *M. abscessus* (Figure 2A-B, Supplemental Figure 2A-B). Two of the antibiotics with the highest accumulation in *M. abscessus* were clofazimine and bedaquiline (Figure 2A), consistent with previous measurements^18^. Concordant with these high uptake values, both clofazimine and bedaquiline display high potency against *M. abscessus*^23^. However, aside from these compounds, the distribution of accumulation of antibiotics is shifted lower compared to the entire drug library (Figure 2C-D). This result is driven in part by the apparent lack of accumulation of penicillin and cephalosporin antibiotics (Figure 2B), which are effectively degraded by *M. abscessus*^24^ and therefore not measured after incubation. Notably, delamanid and thiostrepton were the only two antibiotics aside from clofazimine and bedaquiline that exhibited accumulation within the top 10% of compounds in the library (Figure 2A). Delamanid and related nitroimidazole antibiotics do not display activity against *M. abscessus* likely due to an inability to activate these prodrugs^25,26^, but thiostrepton is a potent inhibitor of *M. abscessus* growth^27^. The relative success of these highly accumulating antibiotics reinforces the notion that identifying antibiotic candidates that effectively penetrate bacteria might result in more successful treatments. Further, the general lack of antibiotic representation amongst highly accumulating compounds suggests that there is substantial room for improvement in the design of drugs that will penetrate mycobacteria.

**Figure 2.**
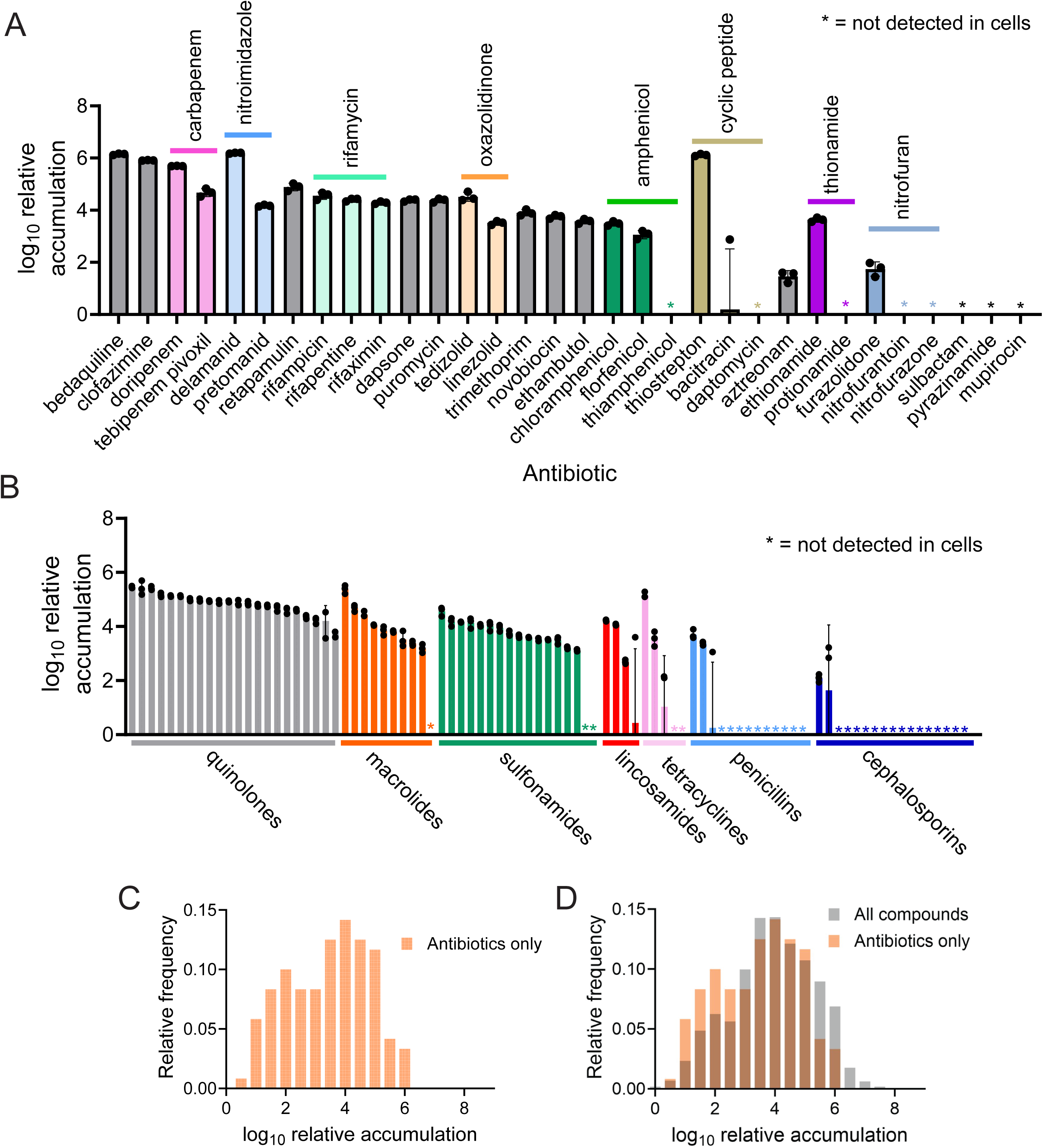
Antibiotics generally display low accumulation in *M. abscessus*. **(A)** LC-MS measurement of relative accumulation of antibiotics of indicated classes. Data represent mean +/- SD. n=3 independent cultures. **(B)** LC-MS measurement of relative accumulation of indicated antibiotics. For antibiotics in a class with more than one member, antibiotic class is indicated above data. Data represent mean +/- SD. n=3 independent cultures. **(C)** Histogram displaying distribution of relative accumulation of antibacterial compounds in *M. abscessus*. **(D)** Overlay of histogram in **(C)** and histogram in Figure 1C.

### High and low accumulating compounds each display structural similarities

Are there chemical moieties that promote uptake that would permit more effective antibiotic design? Of the drugs with the top 10% relative accumulation (Supplemental File 1), there was a significant enrichment of tyrosine kinase inhibitors (Supplemental Figure 2C-D), which commonly display ether- or amine-linked six-membered rings, as demonstrated by ceritinib (Figure 3A). This pattern was also common in other, functionally unrelated compounds in the top 10%, such as pirarubicin (Figure 3A). Phenyl rings connected by linkers several atoms long were also common, as illustrated by the functionally unrelated drugs avobenzone and diminazene (Figure 3A). Also present in the top 10% of compounds were certain steroid-like drugs, such as fluorometholone (Figure 3A).

**Figure 3.**
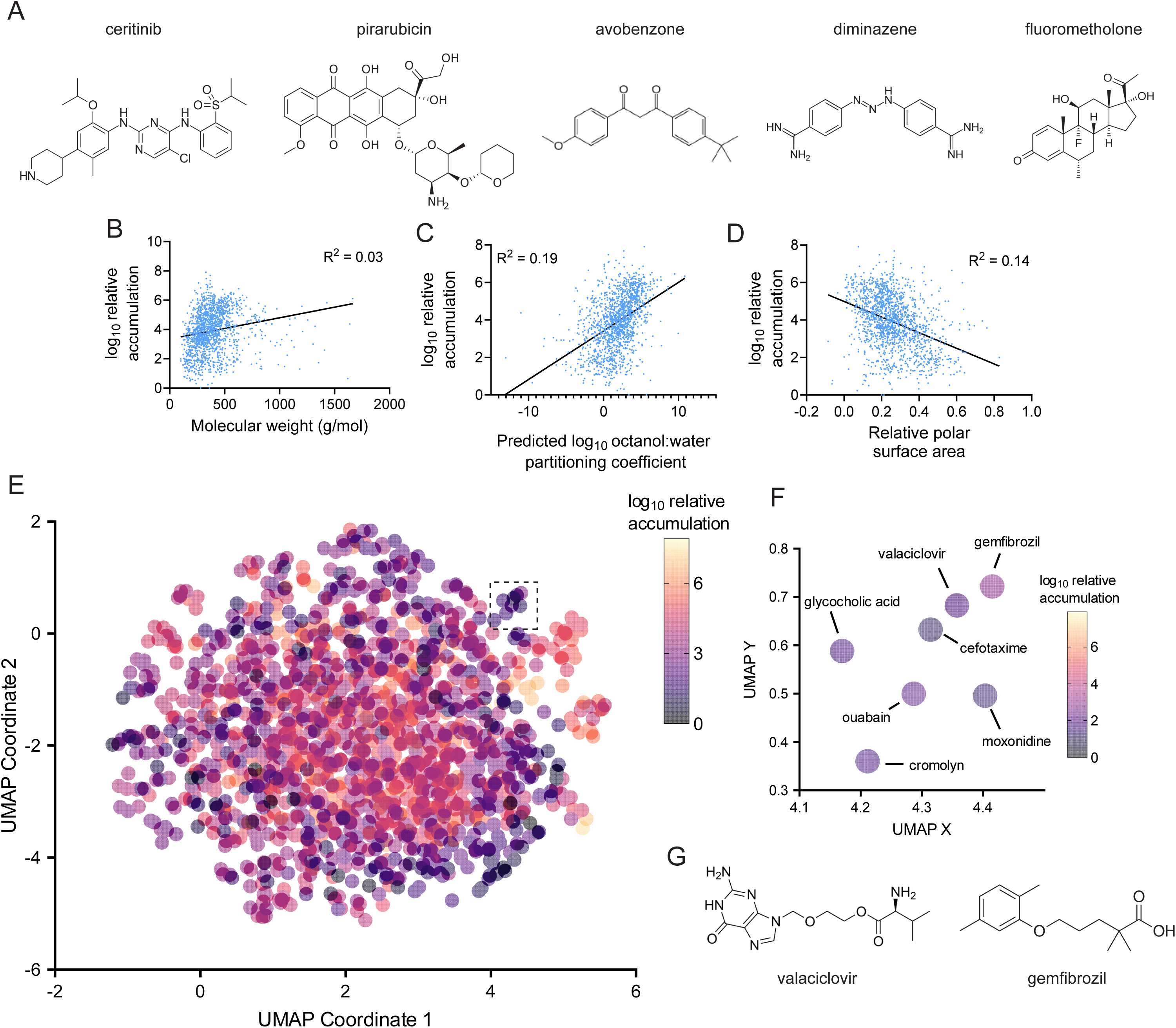
Physical properties do not strongly predict chemical uptake in *M. abscessus*. **(A)** Chemical structures of selected compounds in the top 1% of relative accumulation in *M. abscessus*. **(B)** Correlation between molecular weight **(B)**, predicted octanol:water partitioning coefficient (logP) **(C)**, and relative polar surface area **(D)** with log10 relative accumulation in *M. abscessus*. R^2^ represents the coefficient of determination. **(E)** UMAP depicting fragment-based chemical similarity of 1528 compounds. Log10 relative accumulation for each compound is indicated by color. **(F)** Inset of **(E)** indicated by dotted box. **(G)** Chemical structures of indicated compounds present in **(F)**.

The drugs that accumulated most poorly also displayed some specific patterns. For instance, many contained acetyl groups that could potentially be removed by deacetylases within the cell (Supplemental File 1). Drugs that are metabolized or degraded in this way will not build up in the cell, which would explain their low accumulation values. This class is exemplified by roxatidine acetate (Supplemental Figure 3A), a chemical known to be rapidly deacetylated in humans^28^. The bottom 1% also included compounds such as bisocadyl (Supplemental Figure 3B), which is known to poorly cross the intestinal mucosal barrier^29^, suggesting that some properties of impermeability may be broadly conserved across biology. Interestingly, several of the most poorly accumulating compounds target acetylcholine-binding proteins (Supplemental Figure 3C)^30–32^. Though these compounds do not display any obvious common structural motif, the similarity of their targets suggests that they may have some shared properties that lead to their exclusion from cells.

### General chemical properties are minimally predictive of accumulation

Though examination of these compounds provided some insight into uptake, we sought to identify more generalizable properties of chemicals that could predict their accumulation. We first examined whether any chemical properties commonly used in drug design or previously found to correlate with chemical uptake in other bacteria^14,16^ correlate strongly with drug accumulation in *M. abscessus*. However, neither molecular weight, lipophilicity (as determined by predicted octanol:water partitioning coefficient), relative polar surface area, druglikeness, ring content, or globularity of drugs correlated strongly with accumulation (Figure 3B-D, Supplemental Figure D-G). Notably, this result differs from outcomes in Gram-negative bacteria^14,16^, suggesting that drug uptake is driven by different factors across bacterial genera. Further, these results suggest that drug uptake in *M. abscessus* is likely determined by more nuanced factors than bulk chemical properties.

### Chemical clustering is not consistently predictive of drug accumulation

We next examined whether more complicated structural determinants of drug uptake are captured by chemical clustering using drug fingerprints. This approach compares chemical similarity between compounds based on the presence of specific structural fragments^33^. Similarity between drugs can then be plotted visually through a dimensional reduction that preserves the distance between each drug. We tested whether compounds with high degree of chemical similarity through a fragment-based descriptor, SkelSpheres^34^, displayed comparable levels of accumulation in *M. abscessus*, as that might enable prediction of chemical uptake based on a drug’s location on a chemical similarity plot. However, most compounds are not well-differentiated enough in two-dimensional chemical space to make meaningful predictions (Figure 3E). For the few compounds that did form well-differentiated clusters, the measured accumulation of drugs in those clusters were similar (Figure 3F), and some contained potentially useful information. For instance, fragment-based clustering placed valaciclovir and gemfibrozil, two mechanistically unrelated drugs, adjacent to each other. Structural inspection of these compounds demonstrates their chemical similarity (Figure 3G), and they display similar levels of accumulation in *M. abscessus* (Figure 3F). Thus, fragment-based comparative methods between chemicals may provide insight into drug uptake, but these methods do not discriminate effectively between most drug-like chemicals, rendering their general predictive power low.

### Transfer learning-based methods enable binary prediction of drug accumulation

Given the relatively low predictive power of chemical properties and compound clustering on drug accumulation, we sought to identify a method that could recognize more complex patterns that dictate chemical uptake. To that end, we applied a deep learning approach^35^ that utilizes transfer learning, as our LC-MS dataset is too small by itself to develop a new model through traditional deep learning approaches. However, transfer learning-based methods show promise even when trained on datasets as small as several hundred data points^35^, suggesting that the 1528 compounds measured in this work might be enough to successfully train a model. We first attempted to produce a classifier that could produce a binary prediction as to whether a given chemical structure represented by a textual Simplified Molecular Input Line Entry System (SMILES) string is likely to accumulate in *M. abscessus* or not (Figure 4A). Two different classifiers were generated: one, for which positive accumulation is defined as an accumulation value that would reside within the top 50% of values in Figure 1D, and a second, for which positive accumulation corresponds to a top 20% accumulation value. Each model was trained on 80% of the LC-MS measurements in Figure 1D, with 10% of the compounds used as a validation set and 10% of compounds removed from the training process to serve as a test set for the finalized model (Figure 4A). After training the top 50% classifier, the finalized model was used to predict accumulation of the test compounds, and the results were analyzed using a receiver operating characteristic (ROC) curve (Figure 4B). The area under the ROC curve for this model was 0.90, indicating that the model was highly accurate in predicting drug accumulation in the test set. The second classifier that was trained based on compounds with top twentieth percentile accumulation was highly predictive, with an area under the ROC curve of 0.86 (Figure 4C). Together, these results demonstrate that deep learning-based methods can accurately predict drug accumulation in *M. abscessus*, suggesting that this approach can potentially predict chemicals that will display high uptake into mycobacteria. More broadly, the fact that these models can consistently predict chemical uptake indicates that there are highly regular rules that underlie chemical permeability.

**Figure 4.**
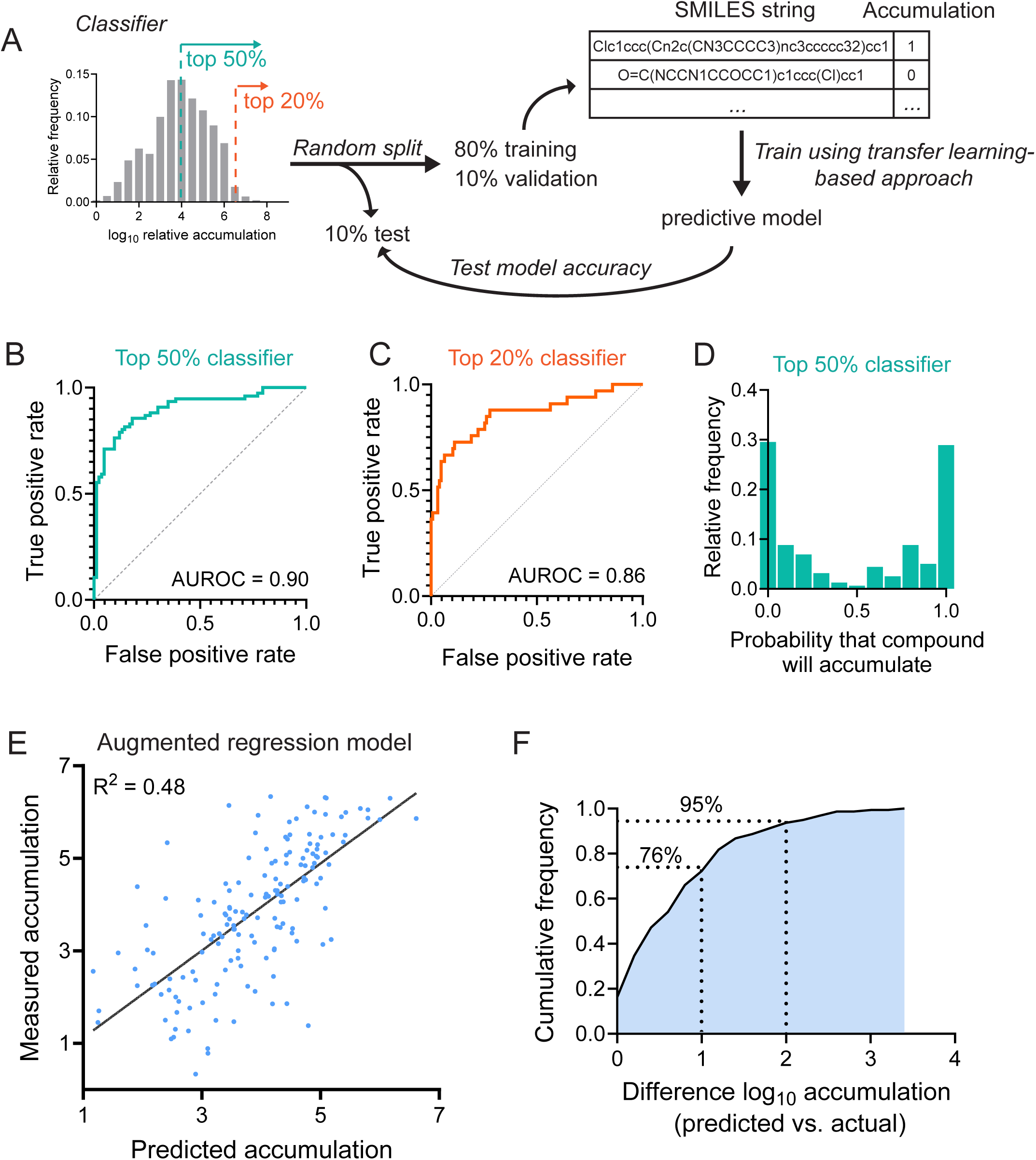
Transfer learning-based models accurately predict chemical accumulation. **(A)** Schematic of classifier-based deep learning approach to predict chemical accumulation. **(B)** Receiver operating characteristic (ROC) curve for a classifier that considers a top 50% relative accumulation value to correspond to accumulation. **(C)** Receiver operating characteristic (ROC) curve for a classifier that considers a top 20% relative accumulation value to correspond to accumulation. **(D)** Histogram of predictions made by the top 50% classifier in **(B)** on the corresponding test set. **(E)** Correlation of predicted and measured log10 relative accumulation of test set compounds by an augmented regression model. R^2^ represents the coefficient of determination. **(F)** Cumulative distribution function for the difference between predicted and actual log10 relative accumulations in **(E)**. AUROC: area under receiver operating characteristic curve.

### Regression-based deep learning models are predictive of drug accumulation

Classifier-based models of drug accumulation produced highly accurate predictions; however, these models provided a bimodal set of predictions, with most compounds having probabilities of accumulation either very close to 0 or very close to 1 (Figure 4D). This is likely a result of the fact that any compound that is not close to the critical threshold of the fiftieth percentile will be likely to have a very high or low probability of accumulation. To provide more quantitative predictions for compounds across the spectrum of accumulation, we used a regression-based predictive model, which is trained directly on the values of relative accumulation for each compound (Figure 1B). This regression-based approach produced a model that was able to predict values for drug accumulation in the test set (Figure 4E), with the majority of compounds having predicted accumulation that fell within one log10 of the actual value (Figure 4F). These results suggest that regression-based models can produce relatively accurate predictions of chemical accumulation, which might enable effective rank-ordering of compounds based on their expected accumulation.

### Predictive models are mildly sensitive to changes in training method

To both gain insight into the robustness of these models and to identify improvements to their predictive power, the effects of several changes to training method on prediction accuracy were tested. First, classifier and regression models were trained using different random splits of the LC-MS data, resulting in unique training, validation, and test sets. These alterations resulted in moderate shifts in the accuracy of both classifier (Supplemental Figure 4A) and regression (Supplemental Figure 4B) models, suggesting that the specific composition of each data set can influence how accurate the final model is. This indicates that a larger training set might further increase the accuracy of these models by reducing reliance on specific compounds being present in the training and validation sets.

Next, classifier and regression models were tested with augmentation of SMILES inputs. For one chemical structure, there exist many unique SMILES strings that can represent that structure. The predictive models produced in this work are trained on the specific SMILES strings used to represent each chemical, which can produce an artificial bias in the model towards those specific SMILES strings rather than the underlying chemical structures. As a result, models of this type can sometimes be improved by training on data sets that use multiple SMILES representations for each compound^36^. SMILES augmentation did not result in improved performance of the classifier model (Supplemental Figure 4C), but augmentation did improve the accuracy of the regression model (Figure 4E, Supplemental Figure 4D). As a result, the augmented regression model (Figure 4E) and the un-augmented classifier (Figure 4B) were used for further experiments.

### Deep learning-based models accurately predict accumulation of quinolone antibiotics

To validate these predictions, we simulated a drug development process by testing structural variants of specific compounds. We used the quinolone class antibiotics, which are compounds that share a quinolone structural core and interfere with DNA gyrase and topoisomerase activity^37^. Eight quinolone antibiotics that were not present in our original library were chosen, and their uptake was predicted using both the classifier model and the augmented regression model (Figure 5A). Predicted accumulation of these compounds by the regression model spanned a 100-fold range, and classifier predictions of the probability of drug accumulation spanned a range from zero to one (Figure 5A). Notably, the regression and classifier predictions were broadly in agreement, with some minor discrepancies. LC-MS measurement of accumulation of these quinolones (Figure 5B) demonstrated a high degree of correlation between measured accumulation and both the regression predictions (Figure 5C) and classifier predictions (Figure 5D), demonstrating that this method is capable of determining chemical accumulation levels of structurally related compounds. Unfortunately, none of the quinolone antibiotics tested display any activity against *M. abscessus* (Supplemental Figure 5A), likely reflecting a lack of enzymatic inhibition due to the variable and species-specific nature of quinolone binding sites^38,39^. These results highlight that the structural changes that promote drug activity are not necessarily coupled to changes that promote uptake. Consequently, pre- emptive identification of chemical variants that accumulate effectively might substantially streamline the process of drug optimization by focusing efforts only on compounds that are both predicted to accumulate and that display effective enzymatic inhibition.

**Figure 5.**
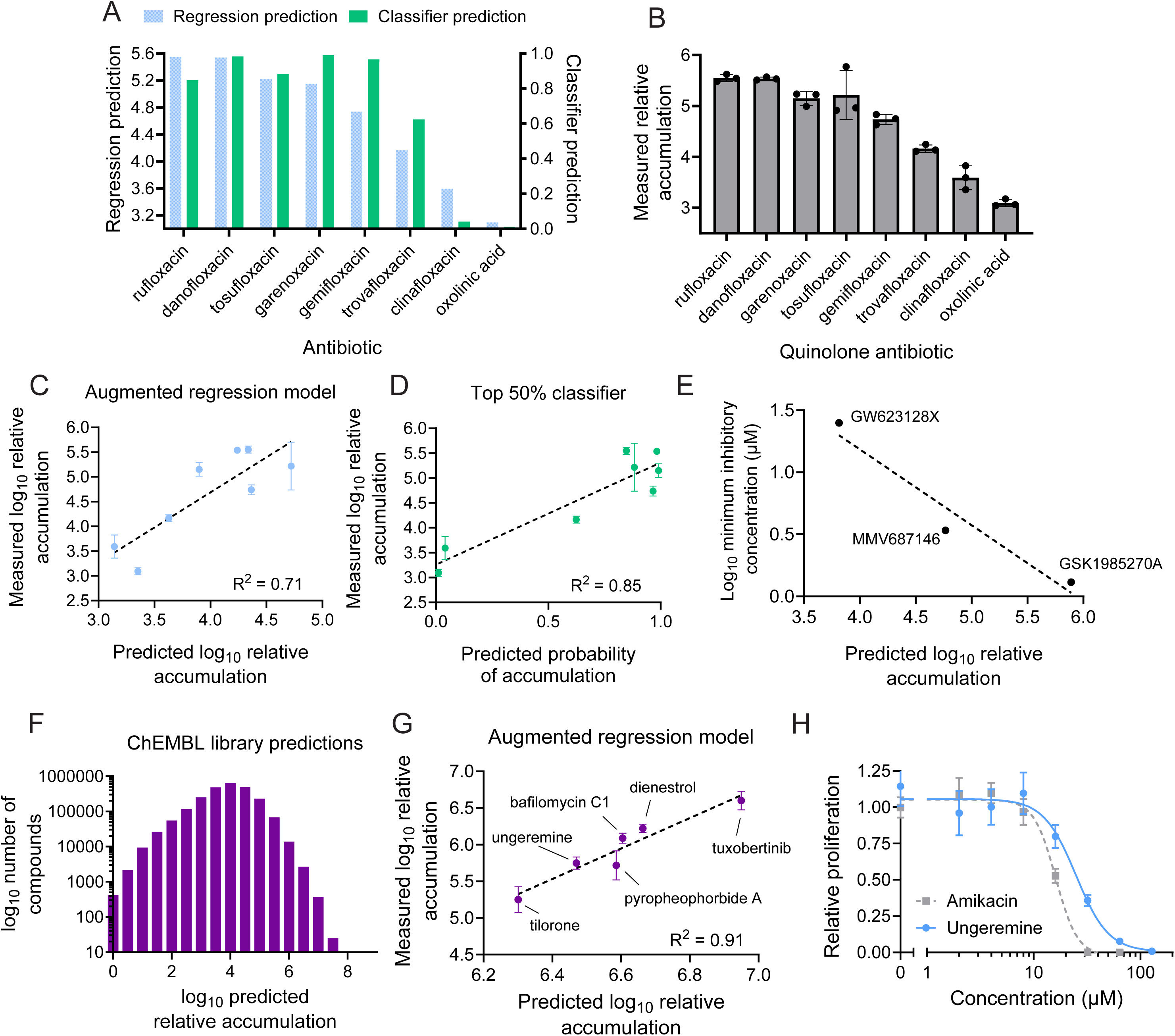
Prediction of chemical uptake reveals compounds with high accumulation. **(A)** Prediction of chemical accumulation for 8 quinolone antibiotics not previously measured by both the augmented regression model in Figure 4E and the classifier in Figure 4B. **(B)** LC-MS measurement of peak area of cell-associated fraction of indicated quinolone antibiotics in *M. abscessus* normalized to peak area of each compound in the media used for the experiment prior to incubation with bacteria. Relative accumulation is normalized to the compound with the lowest accumulation in Figure 1B. Data represent mean +/- SD. n=3 independent cultures for each chemical compound. **(C)** Correlation between the augmented regression model predictions in **(A)** with the LC-MS measurements in **(B)**. Each dot represents one quinolone antibiotic. Data represent mean +/- SD. n=3 independent cultures for each chemical compound. **(D)** Correlation between the classifier model predictions in **(A)** with the LC-MS measurements in **(B)**. Each dot represents one quinolone antibiotic. Data represent mean +/- SD. n=3 independent cultures for each chemical compound. **(E)** Correlation between augmented regression model predictions of uptake of the indicated three MmpL3 inhibitors with log10 minimal inhibitory concentrations for each drug against *M. abscessus*. Log10 minimal inhibitory concentrations were not measured in this study, but instead calculated based on previous work^40^. **(F)** Histogram of the distribution of predicted log10 relative accumulation of 2.4 million compounds in the ChEMBL library. Bins represent log10 number of compounds in each range of predicted accumulation. **(G)** Correlation between augmented regression model predictions of uptake of the indicated six compounds with LC-MS measurements of the relative accumulation of those six compounds. Data represent mean +/- SD. n=3 independent cultures for each chemical compound. **(H)** Relative proliferation of *M. abscessus* as measured using an autoluminescent strain in the presence of the indicated concentrations of ungeremine or amikacin. Proliferation is normalized to vehicle-treated condition. Data represent mean +/- SD. n=6 independent cultures for each drug dose.

### Predicted accumulation correlates with efficacy of compounds with known activity against *M. abscessus*

To test whether predicted chemical accumulation is a useful indicator of the efficacy of drug candidates, we chose to examine a set of three compounds in development (Supplemental Figure 5B) that have all been validated to inhibit *M. abscessus* MmpL3^40–42^, which is a transporter essential for mycomembrane synthesis^43^. Despite targeting the same protein, these compounds display substantially different potencies against *M. abscessus*. Since each of these compounds inhibits the same target in *M. abscessus*, we hypothesized that uptake might explain the differential efficacy of these three compounds. Indeed, each compound exhibited a unique predicted accumulation, and these predictions were strongly anti-correlated with the minimal inhibitory concentration of each drug (Figure 5E). Together, these results suggest that prediction of chemical uptake can be used to pre-emptively identify more effective drug candidates.

### Prediction of accumulation of 2.5 million chemicals identifies high-uptake compounds

Rather than starting from a known antibiotic, many drug development efforts focus on finding new chemical scaffolds with moderate activity against a target or against live bacteria that can be subsequently optimized^44^. We reasoned that prediction of chemical scaffolds with high accumulation might aid in identifying compounds that should be prioritized during this drug development process. To this end, we predicted the accumulation of 2.5 million compounds in the ChEMBL chemical library^45^ using the augmented regression model (Figure 5F). These predictions resulted in a similar distribution of predicted accumulation compared to our measurement of accumulation of 1528 compounds (Figure 1D). Notably, this analysis identified thousands of compounds with predicted accumulation that would have fallen in the top 20% of the 1528 compound drug library, suggesting that those chemicals could represent scaffolds that would accumulate in *M. abscessus*. Six compounds that have never been tested in *M. abscessus* were chosen to validate the accuracy of these predictions; these chemicals were selected on the criteria of high predicted accumulation, LC-MS compatibility, and practical availability, since most compounds in the ChEMBL library are not readily commercially available. LC-MS measurement of these six compounds demonstrated substantial correlation between predicted and measured accumulation (Figure 5G), indicating that these predictive methods are effective across a wide range of drug-like chemicals. Further, these predictions establish a large number of chemical structures that are likely to accumulate in *M. abscessus* and therefore might serve as promising scaffolds for future drug development efforts.

### Identification of high accumulating compounds that display toxicity to *M. abscessus*

Given the high level of accumulation observed for these six compounds in *M. abscessus*, we questioned whether any of these chemicals display toxicity to the bacterium. Indeed, one compound, pyropheophorbide a, displayed modest toxicity to *M. abscessus*, though it failed to fully prevent bacterial proliferation (Supplemental Figure 5C). Pyropheophorbide a is a photosensitizer that produces reactive metabolites upon exposure to light^46^. Its toxicity even in the absence of active light stimulation (Supplemental Figure 5C) suggests that *M. abscessus* might be particularly sensitive to the reactive species produced by pyropheophorbide a. This indicates that pyropheophorbide a might represent a promising scaffold for development of photosensitizers that could be used to treat cutaneous infections caused by *M. abscessus* or other mycobacteria (such as *Mycobacterium marinum and Mycobacterium ulcerans)*, as photodynamic therapy has shown promise against a variety of mycobacteria^47^.

In addition to pyropheophorbide a, a second high-accumulating compound, ungeremine, inhibited growth with potency similar to the widely used standard-of-care antibiotic, amikacin (Figure 5H)^48^. Ungeremine displays toxicity to several bacterial species^49^ as well as to certain cancer cell lines by inhibiting topoisomerase enzymes^50,51^, and it is tolerated with minimal body weight loss in mice^51^, suggesting that ungeremine derivatives might be clinically viable. Further, though ungeremine likely targets similar processes as the highly successful quinolone antibiotics, it bears little structural resemblance to quinolones (Supplemental Figure 5D), suggesting that it may represent a promising orthogonal scaffold for optimization in *M. abscessus*, where clinically used quinolones display low uptake^18^ and are therefore not highly effective.

## Discussion

Identification of promising drug candidates is challenging. Drugs must maintain properties that allow them to be effectively distributed through the human body, accumulate to a high level in the cell of interest, and ultimately engage their enzymatic target. None of these qualities alone is sufficient to make a useful drug; however, bypassing experimental screening for any of these criteria could vastly improve the efficiency of drug development. This work demonstrates that chemical permeability can be effectively measured on a large scale in a pathogenic mycobacterium using LC-MS, and that these measurements can be used to build predictive models that allow for pre-emptive identification of chemicals that will display high intrabacterial accumulation. We propose that this approach could enable more streamlined drug discovery that focuses increased attention on compounds with high predicted accumulation. This increased efficiency could help make it more plausible to develop drugs targeted at rarer pathogens like *M. abscessus*, which typically are relegated to being treated with repurposed drugs that have do not necessarily have optimal activity or uptake. Furthermore, future measurement of chemical accumulation in additional cell types, from other pathogens to human cells, could enable development of a wide range of predictive models to produce more selective and effective accumulation of drugs in targeted cell types.

What are the rules that determine chemical uptake? Our results suggest that chemical accumulation is perhaps not driven by the presence or absence of certain functional groups, as the most prolific accumulators (Figure 3A) display reasonable chemical diversity. Instead, uptake may be driven by structural motifs that are less rigidly defined than by a single functional group. For instance, the structures of avobenzone and diminazene (Figure 3A), two highly accumulating compounds, display broad similarities. The only functional groups that they share in common are phenyl rings; yet, the shapes of the compounds are similar and both exhibit high levels of accumulation. These types of patterns are not easily captured by searching for enrichment of functional groups or even by general chemical properties, suggesting that more complex pattern-searching methods like those enabled by deep learning are critical to detect the chemical motifs that determine drug uptake.

A better understanding of the principles that drive drug uptake might enable development of highly selective therapies that will only accumulate to a significant degree in certain cells. For antibiotics, designing drugs that accumulate highly in pathogenic bacteria but not in host cells or in members of the microbiome could reduce drug side effects and increase the ability to treat with higher doses of antibiotic. However, that possibility is predicated on the assumption that different organisms or cell types display unique accumulation patterns. This work suggests that this assumption may be accurate, as comparing our results to prior work in Gram-negative bacteria^14,16^ suggests that the patterns that drive chemical accumulation differ dramatically across these divergent bacterial species. Our work focuses on *M. abscessus*, a highly drug-resistant organism that is likely among the most effective cell types at excluding chemicals. It is conceivable that mammalian cells, lacking the barriers provided by a cell envelope, may simply take up all drugs to a higher degree than *M. abscessus*, making it more difficult to identify compounds that selectively accumulate in the pathogen and not the host. Measurement of drug accumulation across these diverse cell types will inform the degree to which prediction of chemical uptake can result in selective therapies.

We focused primarily on the chemical determinants that lead to drugs being taken up or excluded from cells. However, the mechanisms that drive those patterns are fundamentally produced by the biology of the bacterium, including the barriers posed by the cell envelope, active drug efflux, and drug metabolism. As a result, chemical uptake is not a fixed property of bacterial cells. Drug efflux and modification rates are determined by the metabolic state of the bacterium, as well as by transcriptional programs that respond to the presence of drugs. These biological processes may differ depending on the physiological contexts in which the bacteria are placed. Moreover, these properties might vary between different strains even within a species. Future efforts to measure and predict drug uptake of pathogens in the context of infection and drug treatment will be critical to account for these variations. In addition to adding complexity to prediction of uptake, these biological processes present a potential therapeutic target. Applying the methodologies outlined in this work to cells undergoing inhibition of specific efflux pumps or drug metabolizing enzymes could broadly reveal the substrates of those enzymes. These insights could bolster ongoing efforts to inhibit bacterial drug-resistance pathways^52^ and could result in rational combination of drugs with efflux inhibitors, which might improve the effectiveness of antibiotics for clinically challenging organisms like *M. abscessus*. Together, the prediction of chemical uptake has the potential to positively impact drug development, and future work to expand these predictions into other contexts could have a far-reaching impact on the process of drug development and on our understanding of the biology that underlies selective cellular permeability to chemicals.

## Acknowledgements

We thank the Harvard Center for Mass Spectrometry for assistance with LC-MS experiments and thank Kritee Mehdiratta, Sam Zinga, Lauren Sullivan, and all members of the Rubin lab for thoughtful feedback on this work. M.R.S. received support as a Merck Fellow of the Damon Runyon Cancer Research Foundation, DRG-2415-20. E.J.R. was supported by NIH/NIAID under award number R01AI179642 and by the Bill & Melinda Gates Foundation (INV-080847).

## Materials Availability

All reagents generated in this study are available upon request from the corresponding author.

## Data Availability

All relevant data generated in this study are present within the manuscript and Supplemental Information.

## Code Availability

All code utilized in this study, along with all deep learning predictive models are available upon request. A web interface to query these predictive models will be made available prior to final publication.

## Materials and methods

### Strains

All experiments were performed in the *Mycobacterium abscessus subspecies abscessus* type strain (ATCC19977).

### Mycobacterial culturing conditions

*M. abscessus* liquid cultures were grown in Middlebrook 7H9 broth (271310, BD Diagnostics) with 0.2% (v/v) glycerol (GX0185, Supelco), 0.05% (v/v) Tween-80 (P1754, MilliporeSigma), and 10% (v/v) oleic acid-albumin-dextrose-catalase (OADC) (90000- 614, VWR) or 10% (v/v) albumin-dextrose-catalase (ADC) composed of 50 g/L albumin, 0.03 g/L catalase, 8.5 g/L NaCl, and 20 g/L dextrose. Cultures were shaken at 150 r.p.m. at 37°C.

### Measurement of chemical accumulation

96-well plates were prepared containing 250 µL of 7H9 + 0.5% (v/v) glycerol + 10% (v/v) ADC with 40 µM of a single drug from the DiscoveryProbe FDA-Approved Drug Library (L1021, APExBIO) in each well. Plates were stored at -80°C until use. After thawing, 50 µL of media was transferred from each well to 3 unique plates to be used for experiments. 50 µL of media from each well of the original plate was then combined to measure initial drug concentrations in media. Combined media was mixed by vortexing and stored at - 80°C until extraction.

*M. abscessus* type strain (ATCC19977) was grown until mid-log phase (OD600 of 0.6-0.8) in 7H9 + 0.5% (v/v) glycerol + 10% (v/v) ADC. Cultures were pelleted at 3200 x *g* for 10 minutes at room temperature then resuspended in 7H9 + 0.5% (v/v) glycerol + 10% (v/v) ADC at a final OD600 of 3. Cells were added in biological triplicate to pre-prepared 96-well plates with 7H9 + 0.5% (v/v) glycerol + 10% (v/v) ADC containing a final concentration of 20 µM of drug in each well resulting in a final OD600 = 1.5. Cells were incubated with drugs for 4 hr at 37°C with shaking at 150 rpm. After incubation, cultures were combined in a conical tube, then pelleted at 3200 x *g* for 10 minutes at 4°C and washed twice with pre-chilled blood bank saline (89370-096, VWR). Pellets were resuspended in 0.8 mL 3:1:0.004 acetonitrile:methanol:formic acid + 10 nM verapamil (V105, Millipore Sigma) or 10 nM clarithromycin (A3487, Sigma-Aldrich), then transferred to 2 mL tubes containing 0.1 mm silica beads (116911500, MP Biomedicals). Choice of internal standard (verapamil versus clarithromycin) was determined by the exact masses of each compound in a given plate to ensure no overlap between the internal standard and compounds being measured. Bacteria were lysed utilizing a Bead Bug 3 Microtube Homogenizer (D1030, Benchmark Scientific) three times at 45 second intervals at 4000 rpm. Samples were chilled on ice for 2 minutes in between cycles. Samples were pelleted at 21,130 x g for 10 minutes at 4°C. 50 µL of media was extracted with 450 µL 3:1:0.004 (v/v/v) acetonitrile:methanol:formic acid + 10 nM verapamil or 10 nM clarithromycin as an internal standard. Media samples were vortexed for 1 minute at 22°C, then pelleted 10 minutes at 17,000 x g at 4°C. Supernatant from both media and cell pellet extractions was dried using a speedvac concentrator (Eppendorf 5305) 1 hr at 45°C, resuspended in 40 µL 3:1:0.004 (v/v/v) acetonitrile:methanol:formic acid, vortexed 1 minute at 22°C, and pelleted at 17,000 x g at 4°C. 35 µL supernatant was transferred to 9 mm plastic vials (C4000-11, ThermoFisher Scientific) with screw caps (03-376-481, ThermoFisher Scientific) and stored at -80°C until LC-MS analysis.

### Liquid chromatography-mass spectrometry

LC-MS analysis was performed as described previously^18^ using a QExactive+ orbitrap mass spectrometer (ThermoFisher Scientific) with a heated electrospray ionization (HESI) probe, coupled to a Dionex Ultimate 3000 UPLC system. 4 µL of extracted sample was injected into a Kinetex 2.6 µm EVO C18 column (150 x 2.1 mm), with the autosampler and column held at 4°C and 30°C, respectively. The chromatographic gradient consisted of 0.1% formic acid (solvent A) and 0.1% formic acid in acetonitrile (solvent B). The gradient was run as follows: 0-5 min: 1% solvent B, flow rate 0.3 mL/min; 5-15 min: linear gradient from 1-99% solvent B, flow rate 0.3 mL/min; 15-20 min: 99% solvent B, flow rate 0.3 mL/min; 20-25 min: 99% solvent B, flow rate 0.4 mL/min; 25-30 min: 1% solvent B, flow rate 0.3 mL/min. The mass spectrometer was operated in full scan, positive mode.

The MS data acquisition was performed in a range of 100-1500 m/z, with the resolution set to 70,000, the AGC target at 1e6, and the maximum injection time at 50 msec.

After LC-MS analysis, chemical identification was performed with XCalibur 3.0.63 software (ThermoFisher Scientific) using a 5 ppm mass accuracy and a 0.5 min retention time window. Standards were used for assignment of chemical peaks at given m/z for the [M+H]^+^ or [M+2H]^2+^ ion and retention time and were compared to extraction buffer, media alone, and cells alone blanks. Linear range of detection for 1528 compound library was determined by analyzing four dilutions of chemical standards containing 0.875 nmol, 7.875 pmol, 78.75 pmol, and 590 pmol of each chemical. Linear range of detection for compounds measured in Figure 5B and Figure 5F was determined by analyzing four dilutions of chemical standards containing 1 pmol, 10 pmol, 100 pmol, and 1000 pmol of each chemical. Linear range was examined by visual inspection of standard curves as well as the residuals of those standards, and the values used for those checks are listed in Supplemental File 1. Peak areas were normalized to either verapamil or clarithromycin internal standard. Relative chemical accumulation was calculated by normalizing intracellular chemical peak area after 4 hours of incubation to chemical peak area in media prior to incubation with *M. abscessus*.

### Pirarubicin fluorescence measurements

*M. abscessus* was thawed and grown to saturation in 7H9 + 0.5% (v/v) glycerol + 10% (v/v) OADC, then diluted back and grown overnight to an OD600 = 0.5 in 7H9 + 0.5% (v/v) glycerol + 10% (v/v) ADC. A parallel culture was then pelleted 3200 x *g* for 10 minutes and resuspended in 4% paraformaldehyde for 1 hr at 22°C. The fixed culture was then pelleted 3200 x *g* for 10 minutes and resuspended in 7H9 + 0.5% (v/v) glycerol + 10% (v/v) ADC. Fixed and unfixed bacteria were transferred to black 96-well plates (3915, Corning) containing a final concentration of 10 μM pirarubicin (28384, Cayman Chemical). 10 μM pirarubicin was also added to wells containing 7H9 + 0.5% (v/v) glycerol + 10% (v/v) ADC with no cells, as well as to wells containing conditioned 7H9 + 0.5% (v/v) glycerol + 10% (v/v) ADC that was produced by a saturated culture of *M. abscessus* grown for 72 hr. 96-well plates were incubated at 37°C for 160 min with linear shaking at 1000 rpm. Fluorescence was measured with an excitation wavelength of 500 nm and an emission wavelength of 550 nm every 5 minutes.

### Growth curve

Measurement of *M. abscessus* proliferation was performed using an autoluminescent strain previously described^53^. Luminescent *M. abscessus* was thawed and grown to saturation in 7H9 + 0.5% (v/v) glycerol + 10% (v/v) OADC, then diluted back and grown overnight to an OD600 = 0.5–0.8. Cultures were diluted to an OD600 = 0.008 in each well of a white, 96-well plate (655074, Greiner Bio-One) in 200 μL of relevant medium. Pirarubicin, amikacin disulfate (A1774, Sigma-Aldrich), ungeremine (34549, Cayman Chemical), dienestrol (38954, Cayman Chemical), bafilomycin C1 (19625, Cayman Chemical), pyropheophorbide a (21371, Cayman Chemical), tilorone (17868, Cayman Chemical), tuxobertinib (HY-136789, MedChemExpress), rufloxacin hydrochloride (HY- B0902A, MedChemExpress), danofloxacin mesylate (22408, Cayman Chemical), tosufloxacin tosylate (21427, Cayman Chemical), garenoxacin (33599, Cayman Chemical), oxolinic acid (16789, Cayman Chemical), gemifloxacin mesylate (21047, Cayman Chemical), or clinafloxacin hydrochloride (16923, Cayman Chemical) were added at indicated concentrations to media. For all growth curves, plates were sealed with Breathe-Easy membranes (Z380059, MilliporeSigma) and incubated at 37°C without shaking. Luminescence was measured at 0, 24, and 48 hours after inoculation, and relative proliferation was calculated as the ratio of luminescence at 48 hrs to 0 hrs normalized to the vehicle-treated condition. Growth curve data were analyzed using Microsoft Excel 2016 and GraphPad Prism 10.

### Chemical property correlations

Chemical properties (molecular weight, predicted logP, relative polar surface area, druglikeness, globularity, aromatic ring content, and non-aromatic ring content) for each compound were calculated using DataWarrior v06.02.01^34^.

### Chemical similarity clustering

Chemical fingerprints were calculated using SkelSpheres in DataWarrior v06.02.01^34^ and represented as a 2-dimensional UMAP with nearest neighbors set to 100, minimum distance set to 0.5, and Euclidian distance.

### Generation of predictive models

Predictive models were produced using MolPMoFit^35^ in Python v3.9.19, utilizing Pytorch v1.8.1+cpu and Fastai v1.0.61. ChEMBL_1M_SPE pre-trained model^54^ was used as the basis for model training without further fine-tuning, as fine-tuning did not improve performance in prior work^35^. For classifier models, LC-MS measurements of chemical uptake were rank-ordered, then the value of all compounds in the top 50% were set to 1, and the bottom 50% were set to 0. Regression models were trained directly on the log10 relative accumulation value for each compound. To train both classifier and regression models, LC-MS training data were associated with the appropriate SMILES string for each compound as identified by DataWarrior v06.02.01^34^. Data were randomly split 80:10:10, and the same random seeds to split the data were used for classifier and regression models. SMILES augmentation was performed with a target of 25 SMILES variants for training set structures and 5 variants for validation set structures using RDKit v2024.3.5. Models were trained using previously described parameters^35^ over four epochs with a batch size of 128, chunk size of 50,000, and max vocabulary size of 60,000. ChEMBL library^45^ predictions were performed on ChEMBL release 34.

### Statistical analysis

R^2^ values represent the coefficient of determination for indicated linear regressions. All t-tests were unpaired, two-tailed tests. p-values for contingency tables were calculated by chi-squared test.

## Supplemental Figures

**Supplemental Figure 1.**
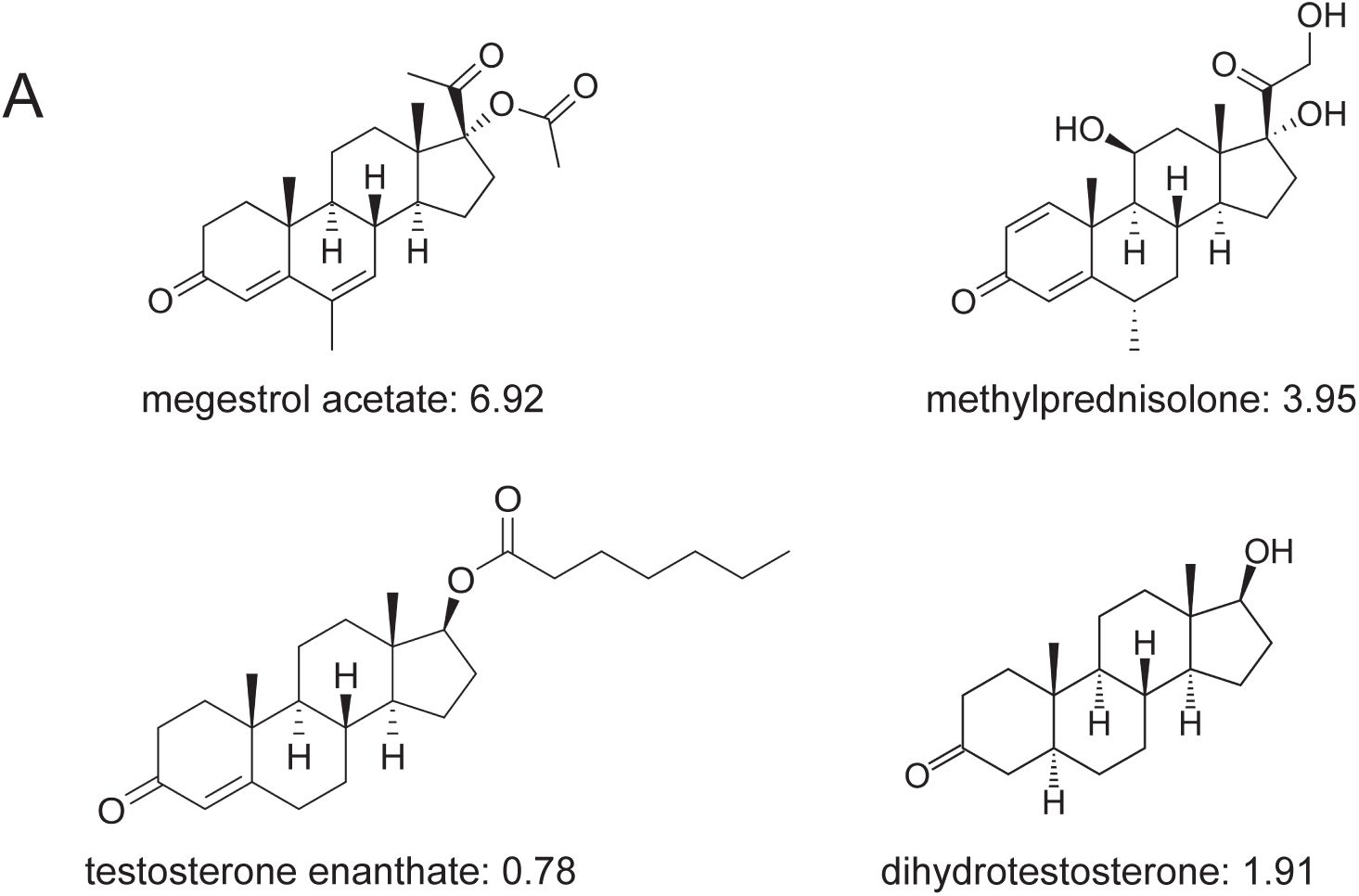
Steroid-like compounds display a wide range of accumulation. **(A)** Chemical structures and log10 relative accumulation of selected steroid-like compounds in *M. abscessus*.

**Supplemental Figure 2.**
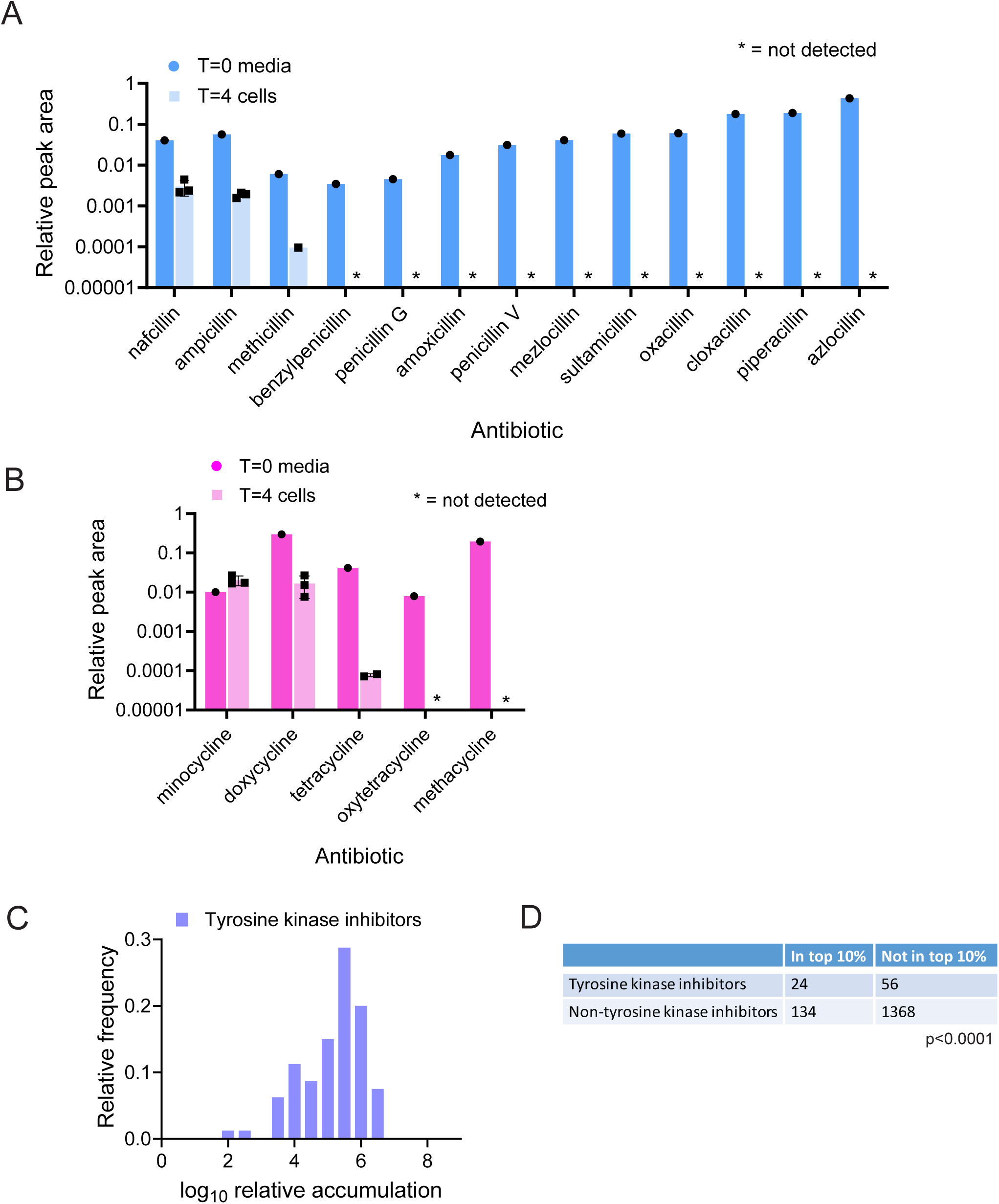
Antibiotics known to be degraded by M. abscessus accumulate poorly. **(A)** Relative peak area in initial media and in cell-associated fraction after 4 hr incubation for the indicated penicillin-related antibiotics. Data represent mean +/- SD. n=3 independent cultures for cell-associated fractions. **(B)** Relative peak area in initial media and in cell-associated fraction after 4 hr incubation for the indicated tetracycline-related antibiotics. Data represent mean +/- SD. n=3 independent cultures for cell-associated fractions. **(C)** Histogram displaying the relative accumulation of tyrosine kinase inhibitors in *M. abscessus*. **(D)** Contingency table categorizing relative accumulation of tyrosine kinase inhibitors versus non-tyrosine kinase inhibitors in M. abscessus. p-value derived from two-sided chi-squared test.

**Supplemental Figure 3.**
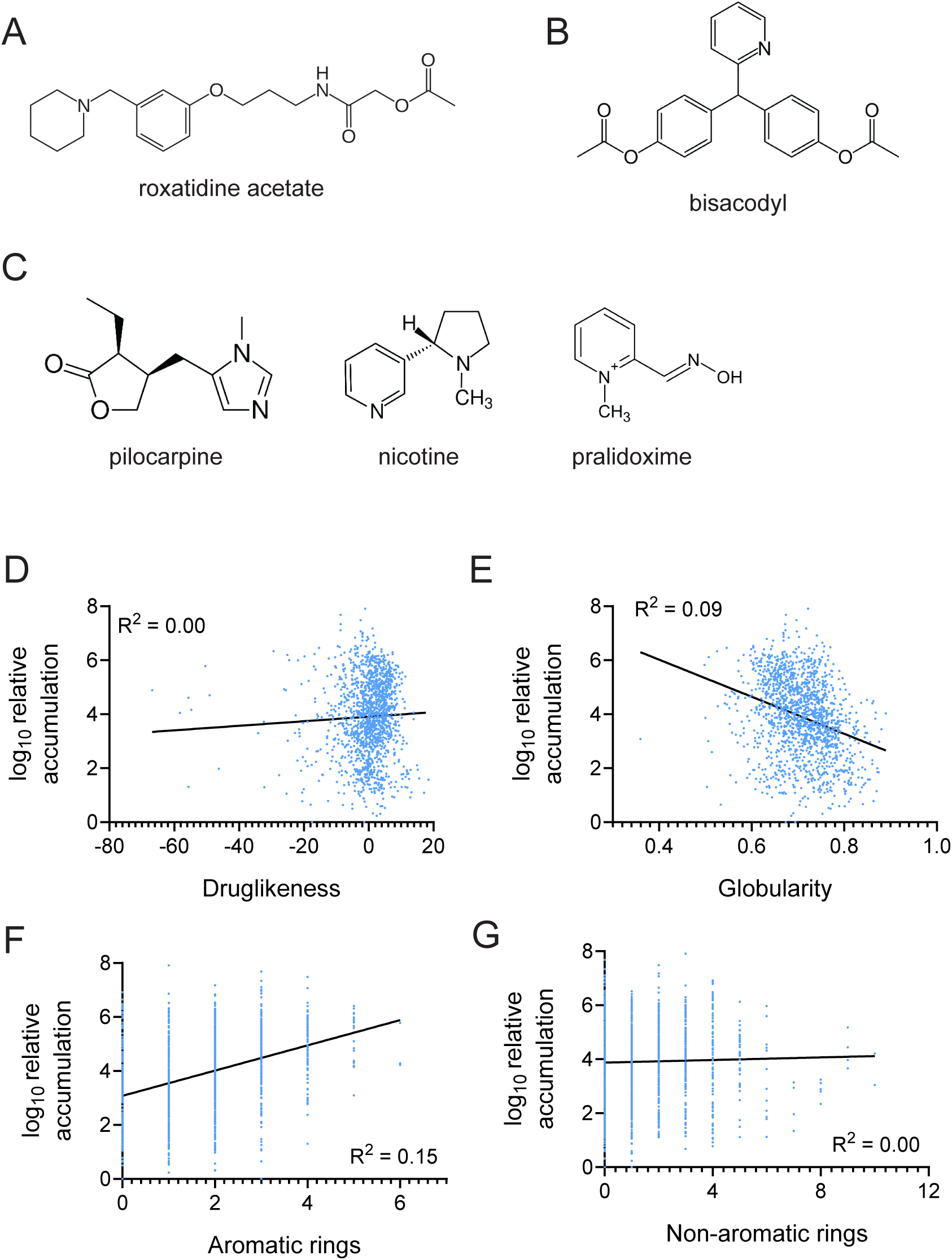
Physical properties poorly predict chemical accumulation in *M. abscessus*. **(A)** Chemical structure of roxatidine acetate. **(B)** Chemical structure of bisacodyl. **(C)** Chemical structures of pilocarpine, nicotine, and pralidoxime. **(D)** Correlation between druglikeness **(D)**, globularity **(E)**, aromatic ring content **(F)**, and non-aromatic ring content **(G)** with log10 relative accumulation in *M. abscessus*. R^2^ represents the coefficient of determination.

**Supplemental Figure 4.**
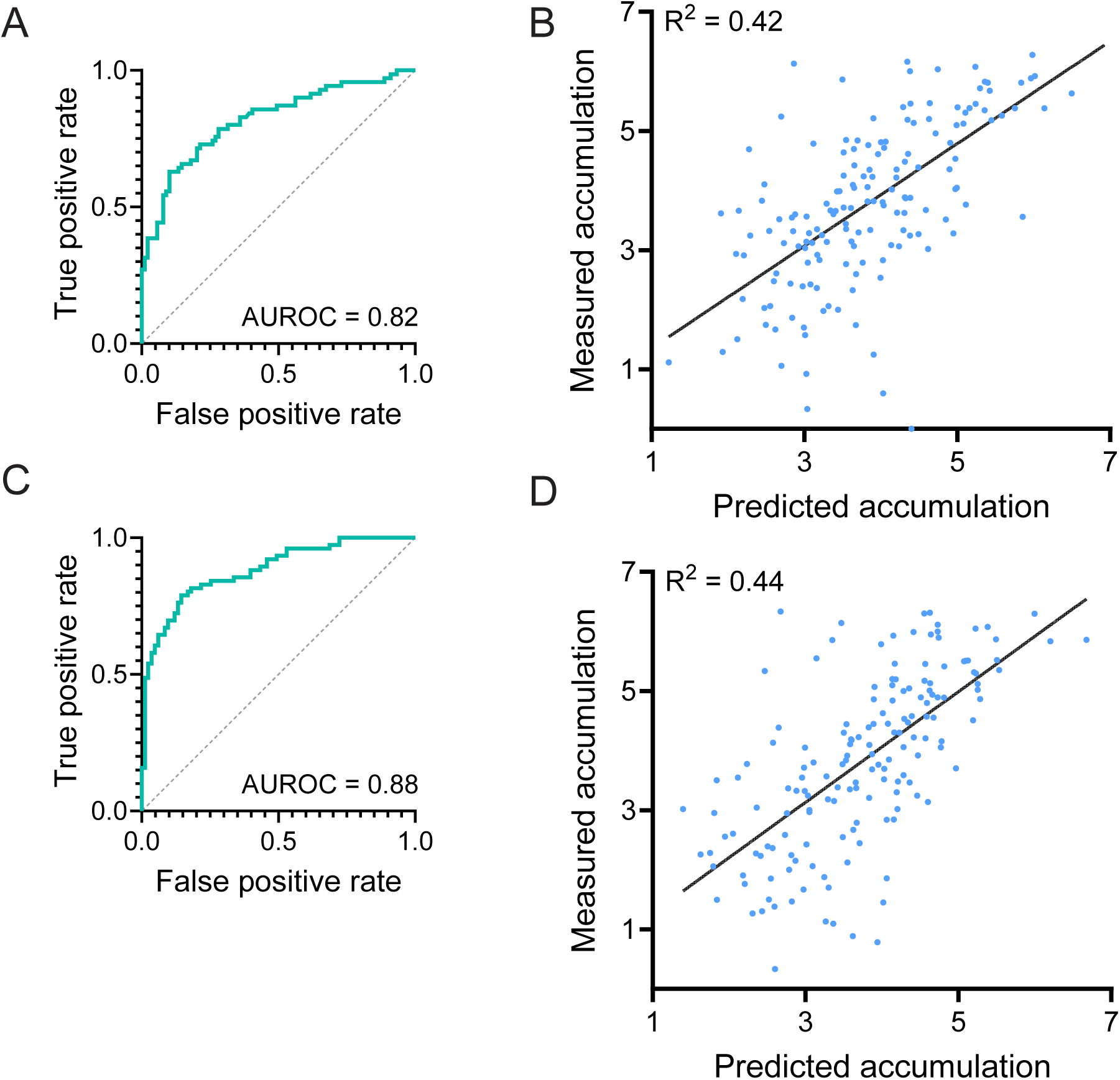
**(A)** Receiver operating characteristic (ROC) curve for a classifier that considers a top 50% relative accumulation value to correspond to accumulation. Classifier was generated using different splitting of training data compared to classifier in Figure 4B. **(B)** Correlation of predicted and measured log10 relative accumulation of test set compounds by an augmented regression model. Regression model was generated using different splitting of training data compared to regression model in Figure 4E. R^2^ represents the coefficient of determination. **(C)** Receiver operating characteristic (ROC) curve for a SMILES-augmented classifier that considers a top 50% relative accumulation value to correspond to accumulation. **(D)** Correlation of predicted and measured log10 relative accumulation of test set compounds by a non-augmented regression model. R^2^ represents the coefficient of determination.

**Supplemental Figure 5.**
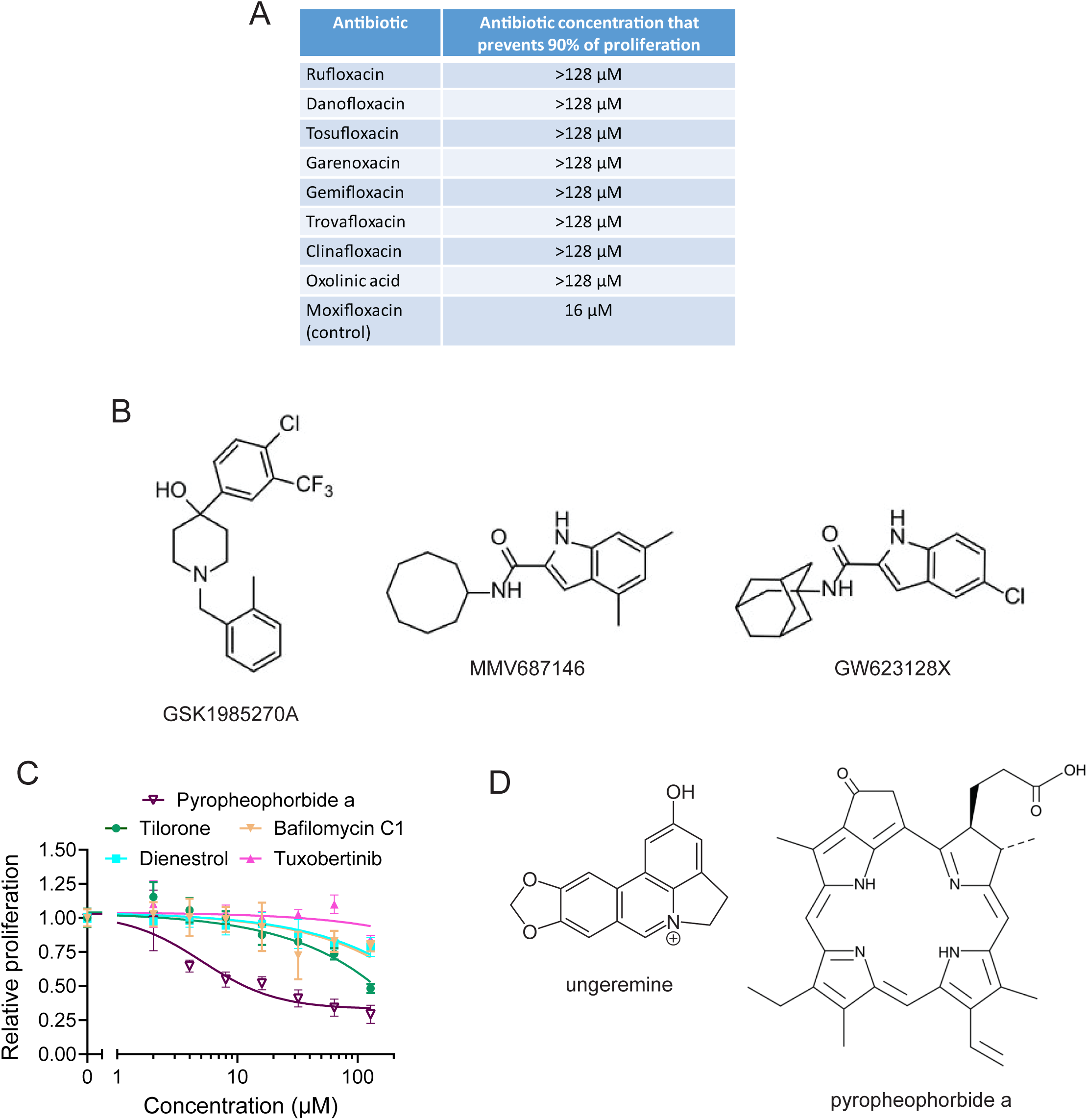
Measurement of antibacterial activity of quinolones and high uptake compounds. **(A)** Antibiotic concentration that inhibits >90% of proliferation of *M. abscessus* as measured by bacterial autoluminescence for 8 quinolone antibiotics measured in Figure 5B as well as clinically used compound moxifloxacin. **(B)** Chemical structures of MmpL3 inhibitors analyzed in Figure 5E. **(C)** Relative proliferation of *M. abscessus* as measured by autoluminescence in the presence of the indicated concentrations of pyropheophorbide a, tilorone, bafilomycin C1, dienestrol, or tuxobertinib. Proliferation is normalized to vehicle-treated condition. Data represent mean +/- SD. n=6 independent cultures for each drug dose. **(D)** Chemical structures of ungeremine and pyropheophorbide a.

